# Twilight length alters growth and flowering time in Arabidopsis via LHY/CCA1

**DOI:** 10.1101/2023.06.09.544111

**Authors:** Devang Mehta, Sabine Scandola, Curtis Kennedy, Christina Lummer, Maria Camila Rodriguez Gallo, Lauren E. Grubb, Maryalle Tan, Enrico Scarpella, R. Glen Uhrig

## Abstract

The plant circadian clock governs growth and development by syncing biological processes to periodic environmental changes. Decades of research has shown how the clock enables plants to respond to two environmental variables that change at different latitudes and over different seasons: photoperiod and temperature. However, a third variable, twilight length, has so far gone unstudied. The duration of twilight changes across the planet, lengthening with latitude and changing across seasons, and yet most circadian experiments are performed in lab environments with no twilight. Here, using controlled growth setups, we show that twilight length impacts plant growth and changes flowering time via the LHY/CCA1 morning module of the Arabidopsis circadian clock. Using a series of progressively truncated no-twilight photoperiods, we also found that plants are more sensitive to twilight length compared to equivalent changes in solely photoperiod. Further, transcriptome and proteome analyses indicated that twilight length alters the regulation of proteins involved in reactive oxygen species metabolism, photosynthesis, and carbon metabolism. Genetic analyses also implicated a twilight sensing pathway from phytochromes D and E, the morning elements of the circadian clock and modulating flowering time through the *GI-FT* pathway. Overall, our findings call for more nuanced models of daylength perception in plants and posit that twilight length is an important determinant of plant growth, development, and circadian function.

## Introduction

The circadian clock is a molecular network that is entrained to a 24-hour oscillatory period by acute changes in various environmental cues such as light and temperature to regulate the diel cycling of physiological activity in all eukaryotes. The plant circadian clock is especially complex, comprised of multiple, interconnected transcription-translation feed-forward loops (TTFLs) that are reset by the onset of dawn and buffered against mild changes in temperature (*1*). The clock itself has far reaching regulatory capacity, directly and indirectly controlling nearly all physiological processes from photosynthesis, energy metabolism, and even biotic stress response (*2*).

Over the lifecycle of a plant, changes in photoperiod and temperature coincide with the circadian cycle to induce developmental changes such as the transition to flowering. These changes are governed by seasonality, which varies depending on the latitude at which a plant is grown. In fact, changes in latitude determine seasonal temperature, photoperiod, as well as the length of dawn and dusk (i.e. twilight length) (*3*). Indeed, the effect of seasonal changes in temperature and photoperiod on both the circadian clock and flowering have been well studied in a range of plant species. For example, mutations in clock genes such as *ELF3* and *PRR7*-orthologs have been linked to the successful migration of bean and alfalfa from tropical to temperate latitudes (and consequently, to the associated changes in temperature and photoperiod) (*4–6*). More recently, a simple translational coincidence model has emerged suggesting that an overlap between clock-controlled transcript phases and light-dependent protein synthesis explains molecular adaptation to new photoperiods due to changing seasons (*7*).

Foundational work on the plant circadian clock has been conducted using rectangular light-dark (LD) cycles with multispectral high-intensity sodium (HPS) or fluorescent tube lights that are turned on or off in a defined time-period. Over the last two decades, this general set-up has led to ground-breaking discoveries in plant chronobiology, from the discovery of the clock components themselves to global transcriptional profiling and modelling under different diurnal light regimes (*7–12*). However, the use of these light setups along with a rectangular LD cycle is a poor match to real-world conditions, particularly at temperate and polar latitudes where relatively long twilights and lower peak light intensities are present. Thus, largely due to technical limitations in laboratory and greenhouse lighting systems, the impact of changing durations of twilight, i.e., the gradual change in light intensity between night and day, remains unstudied in plants.

Given ongoing climate change and the global rise in temperatures (*13*), it is expected that crop varieties and crop species adapted for tropical latitudes will gradually be adopted in temperate zones that feature different seasonal photoperiod changes and twilight length. This phenomenon of northward migration has already been found to occur over the last 40 years in rain-fed wheat, maize, and rice (*14*). Latitudinal migration is also expected to play a role in future agriculture through the expansion of the agricultural climate zone northwards due to global warming (*15*). It is thus crucial to develop a fundamental understanding of how variation in the latitudinal light environment, including changes in twilight length, affects plant growth and development.

Here, we present results from experiments examining the impacts of varying lengths of twilight on the model plant Arabidopsis using multispectral LED lights with precise intensity control. Our results demonstrate that twilight length differentially impacts plant vegetative growth and flowering time. We also show that twilight measurement requires the morning components of the plant circadian clock, and that twilight length alters the internal circadian period. We further present transcriptomic and proteomic data highlighting twilight length-dependent protein-level regulation in select biological pathways. Collectively, our results point to the need for revisiting established ideas regarding core circadian clock functions like photoperiodism by using more natural environmental transitions than previously employed in the field.

## Results and Discussion

To investigate the effects of twilight length on plant growth and development, we devised an experimental set-up that uses a 6-band LED light fixture with fully programmable spectra and sigmoidal ramping of light intensity in controlled growth chambers. We employed this set-up to gain a comprehensive understanding of the impact of changing twilight lengths on *Arabidopsis thaliana* (Col-0) plants at both physiological and molecular levels, studying the effect of a wide range of twilight lengths (0 min., 15 min., 30 min., 60 min., 90 min.) on plant growth. These conditions are approximately comparable to summer twilight lengths at the Equator (∼15 min) up to 59°N (∼90 min), while the 0 min (https://aa.usno.navy.mil/faq/RST_defs) condition is typical for most plant biology experimentation in controlled growth environments and serves as our no-twilight control. Our experiment was designed to isolate the impact of twilight duration on plant growth, separate from other twilight-associated phenomena such as changes in total DLI (daily light integral; the number of photosynthetically active photons delivered over a 24-hour period) and altered wavelength distribution that might impact photosynthetic capacity. Hence, our LD curves ensured that while the length of the dawn/dusk ramp varied, the DLI remained the same between conditions, with relatively minor changes in peak light intensity (<14 μmol difference in peak photosynthetic photon flux density (PPFD) between the various light regimes) (**Fig. 1a; Supplementary Table 1**). We also did not alter the light spectrum over the day as this study focused on the impact of light intensity changes over time, rather than an accurate simulation of natural twilight. However, our light spectrum included a supplemental far-red light component (Red:Far-Red ratio ∼1) according to the recent discoveries by the Imaizumi lab that the Far Red component is important for mimicking flowering under natural conditions (*16, 17*).

**Figure 1:**
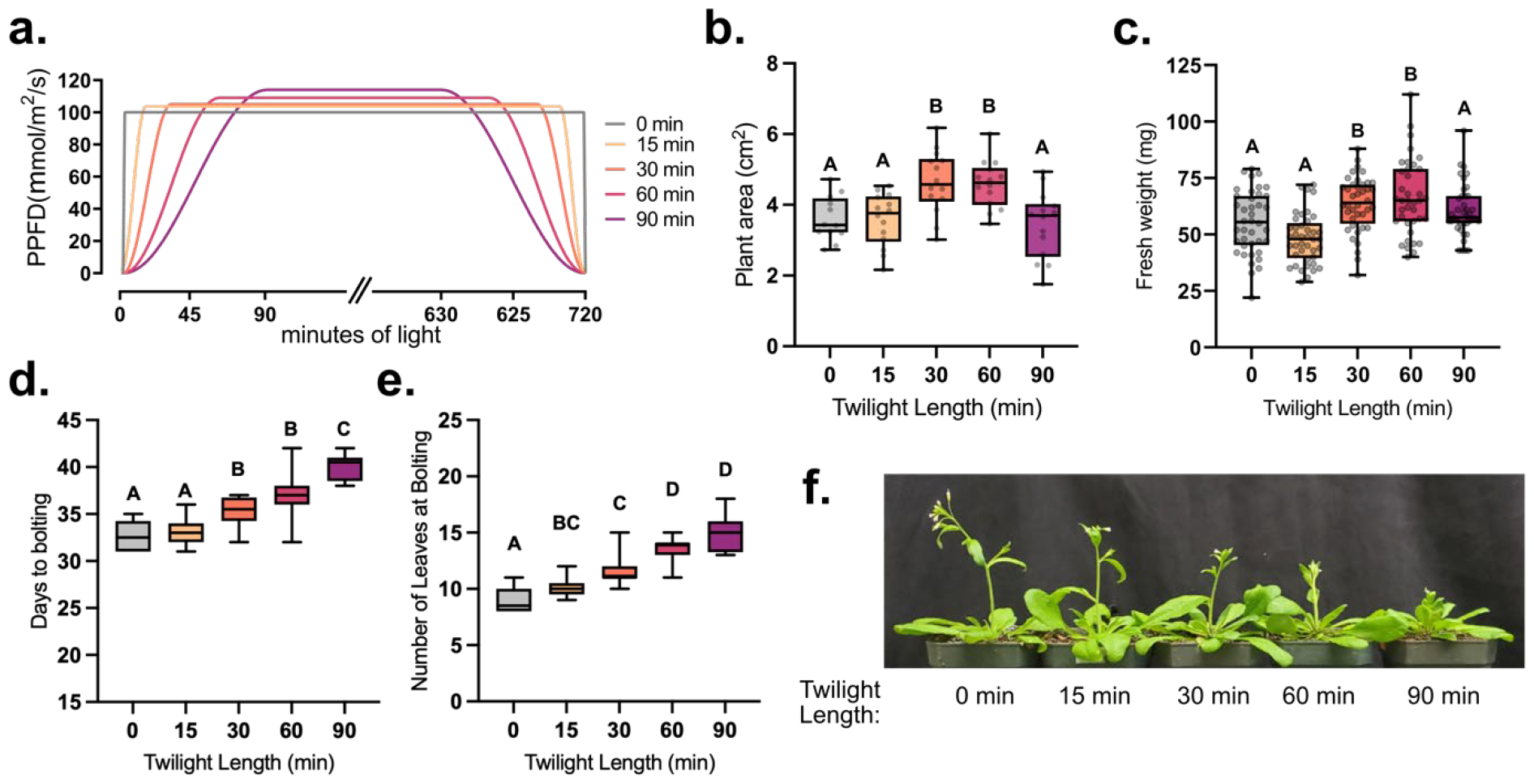
The impact of twilight length on plant growth and development. **(a)** Twilight length experimental schemes. **(b)** Leaf area of wild-type plants as measured using PlantCV from rosette images at 25 days post-imbibition (n>12). **(c)** Fresh weight of plants grown under different lengths of twilight (n>38 plants) measured at 25 days post-imbibition. **(d & e)** Flowering time measured by counting the number of days to bolting and number of leaves at bolting respectively (n>10 plants). **(f)** Representative images of plants at 36 days post imbibition. Letters above all graphs depict significantly different datapoints based on a one-way ANOVA and Tukey’s post-hoc test with adjusted p-value <0.05.

We first analysed plant growth by measuring plant area (total area of the rosette) using overhead RGB cameras and processing images using the open-source PlantCV image analysis software (*18, 19*). Measurements of plant area from images acquired 25 days post imbibition showed that *Arabidopsis thaliana* Col-0 (Arabidopsis) plants grew to larger sizes as twilight length increased to an optimum of 30 and 60 min, with a median increase in plant area of 34% compared to plants grown under a no-twilight rectangular LD cycle (**Supplementary Table 2**). Plants grown under the extended 90 min twilight condition, however, showed no increase in plant size relative to the no-twilight condition (**Fig. 1b**). This trajectory of plant size increasing with longer durations of twilight up to an optimum of 30-60 min was mirrored by plant biomass (fresh weight) measurements taken at 25 days post imbibition. We found that increasing the length of twilight led to a median 16% increase in biomass with a 60-min twilight duration, but no significant change in biomass upon growth with a 90-min twilight compared to the no-twilight control (**Fig. 1c; Supplementary Table 3**). These results were validated in an independent replication of the experiment (**Fig. S1**).

Next, we assessed how this twilight-induced increase in plant size affected plant development by measuring flowering time in wild-type (WT) Arabidopsis plants. Time to flowering or time to bolting can be measured using either direct time measurements (days to bolting) or morphometric indicators such as number of rosette leaves at bolting (*20*). In our twilight length experiment both days to bolting as well as the number of rosette leaves at bolting increased linearly with twilight length, including in the 90 min extended twilight condition (**Fig. 1d & e; Supplementary Table 4**). This result was replicated in an independent experiment (**Fig. S1**). The fact that an extended period of twilight (90 min.) does not result in any increase in vegetative growth (as measured by plant area and biomass) but lengthens flowering time is intriguing as vegetative growth and flowering time are highly correlated in a variety of phenological studies, under different altitudes, latitudes, and genotypes (*21–23*). Indeed, the relationship between vegetative growth and flowering time is regarded as a classic example of a life-history trade-off with strong genetic underpinnings (*24*). In *Arabidopsis thaliana*, previous studies using different photoperiodic regimes have noted a difference in the relationship of flowering time as measured in days to bolting versus leaf count at bolting with decreasing photoperiods. For example, below the so-called “ceiling photoperiod” of ∼8h, flowering time measured in days continues to increase linearly, while leaf number at bolting remains constant (*25*). Thus, our finding that a 90-min period of twilight breaks the relationship between vegetative growth and flowering time calls for a re-evaluation of our understanding of the limits of the growth-reproduction trade-off in plants, particularly at very high latitudes. Further, the fact that this disconnect occurs below the previously defined “ceiling photoperiod” suggests differences between how plants sense photoperiod (or more precisely, how they sense the duration of photosynthetically active radiation) and their means of sensing twilight duration. Indeed, recent targeted research has begun to unravel how photosynthetic daylength is timed differently from “absolute photoperiod” in Arabidopsis under standard experimental LD conditions (*26*).

To further examine the differences between responses to twilight length and photoperiod, we next performed a second experiment where plants were grown under a series of decreasing photoperiods with a rectangular LD scheme and compared to a 30 min. twilight as a control. The photoperiod series was designed to test: (a) how sensitive plants are to minute changes in photoperiod, and (b) whether a minimum amount of light intensity is necessary to result in the plant growth and developmental changes seen under twilight conditions. This second question was guided by previous studies which established that while most plant photoreceptor signaling is saturated at <5 μmol m^−2^ s^−1^, net carbon fixation only occurs beyond 8-40 μmol m^−2^ s^−1^ (*27, 28*). The results described in Figure 1 suggest the hypothesis that as light intensity ramps up and down during twilight, different processes turn on and off at different times, potentially resulting in differential regulation of vegetative growth and flowering time. Hence, under more natural conditions the concept of “photoperiod sensitivity” might in fact encompass two temporally separated phenomena where: (a) plants sense the onset of day and night at very low light intensities (this might also be when the clock is ‘reset’), and (b) metabolic processes more directly tied to growth (e.g., carbon fixation) commence and end at higher light intensity thresholds (potentially in concert with the clock). Prior research into photoperiod sensitivity may not have been able to separate these processes due to the use of rectangular LD cycles.

By contrast, the experimental scheme described in **Fig. 2a** permitted us to test the sensitivity of flowering time to 4, 10, 30, and 60-min reductions in photoperiod. Further, by including a 30-min twilight condition, this experimental design also allowed us to test a light intensity range of 2.6, 7.07, and 41.64 μmol m^−2^ s^−1^ as light detection thresholds that most closely explain the phenotype under a 30-min twilight condition (Light intensity intercepts of the 30-min ramp and successive rectangular photoperiod LD schemes, see **Fig. 2a**.). For instance, if the 5sq condition (a 10-min photoperiod reduction) most closely phenocopied the 30-min twilight treatment, we might conclude that a photoperiodic-light detection threshold close to 7.07 μmol m^−2^ s^−1^ was in effect under this twilight condition. Our results show that only a 30-min reduction in photoperiod (15sq) comes close to the 30-min twilight treatment in terms of a delay in flowering time (**Fig. 2 b & c; Supplementary Table 6**). This implies that if we employ a model where plants purely measure photoperiod (i.e., only the x-axis in **Fig. 2a**) rather than the slope of light intensity at the start and end of day, this measurement must begin and end past ∼40 μmol m^−2^ s^−1^ of light intensity during the ramp to explain the flowering phenotype observed. However, the increase in flowering time over successive reductions in photoperiod is also non-linear, in contrast to the twilight length experiment shown in **Fig. 1 (d & e)**. In addition, a ∼40 μmol m^−2^ s^−1^ light detection threshold is far higher than previously measured thresholds for photoreceptor activation. Collectively, this shows that twilight length impacts flowering time, and that this effect can only partially be explained by differential photoperiod sensing under different twilight length regimes. It is thus imperative that future research into clock function and seasonal sensitivity use more sophisticated experimental designs to account for the differential regulation of growth and flowering under more natural light-dark transitions.

**Figure 2:**
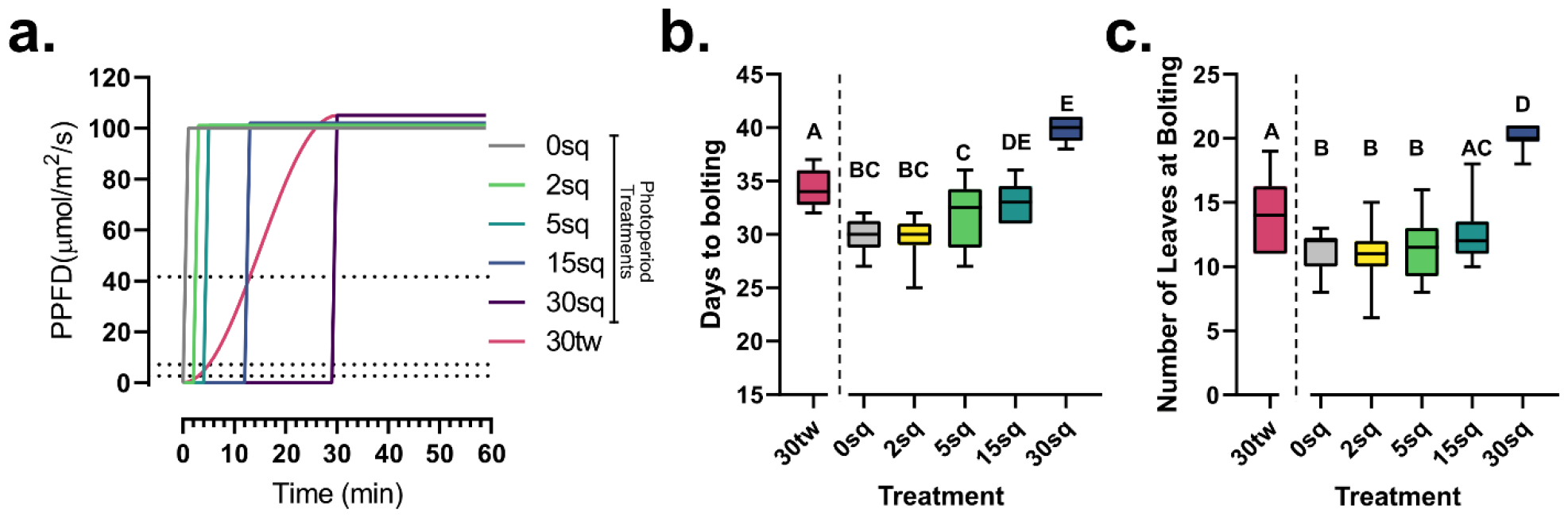
Dissecting the differences between plant responses to twilight and photoperiod. **(a)** Depiction of the light treatments in this experiment from a 12-hour light: 12-hour dark photoperiod (0sq) to 2 min. (2sq), 5 min. (5sq), 15 min. (15sq), 30 min. (30sq) reductions in photoperiod in the morning and evening (only morning shown). A 30-min. twilight condition (30tw) is included as control. Dotted lines depict light intensity at different points along the twilight ramp. **(b & c)** Flowering time measurements in terms of **(b)** days to bolting and **(c)** number of leaves at bolting of wild-type plants grown under the conditions depicted in **(a)**. A minimum of 12 plants per treatment were measured. Letters depict significantly different datapoints based on a one-way ANOVA and Tukey’s post-hoc test with adjusted p-value <0.05.

Since flowering time and seasonal sensitivity are intimately tied to circadian clock function, and because we observed twilight-length associated changes in clock period, we next studied the flowering response of a suite of core clock mutants to different lengths of twilight. We found that both clock activator mutants (*rve 4 6 8*) as well as two clock repressor mutants (*toc1-2, elf4-101*) showed more similar flowering time changes to wild-type plants, with progressive increases in twilight length (**Fig. S2**). However, the double mutant of the morning element of the clock, *lhy cca1* showed a near-complete insensitivity to changes in twilight length, with significant changes in flowering time observed only under the 90-min twilight treatment (**Fig. 3; Supplementary Table 4**). To see if this absence of twilight sensitivity also manifested in growth outcomes, we monitored plant area and biomass for *lhy cca1* plants, finding smaller changes in plant area (**Fig. 3a**) and biomass (**Fig. 3b**) at different twilight lengths compared to wild-type plants (**Fig. 1 a & b**). These results were replicated in an independent experiment (**Fig. S1**). Together with our analysis of twilight and photoperiod, the lack of twilight sensing in the *lhy cca1* mutant suggests that the LHY/CCA1 module of the clock is key to sensing not only effective photoperiod, but also the duration of twilight, i.e., the slope of increasing and decreasing light intensity.

**Figure 3:**
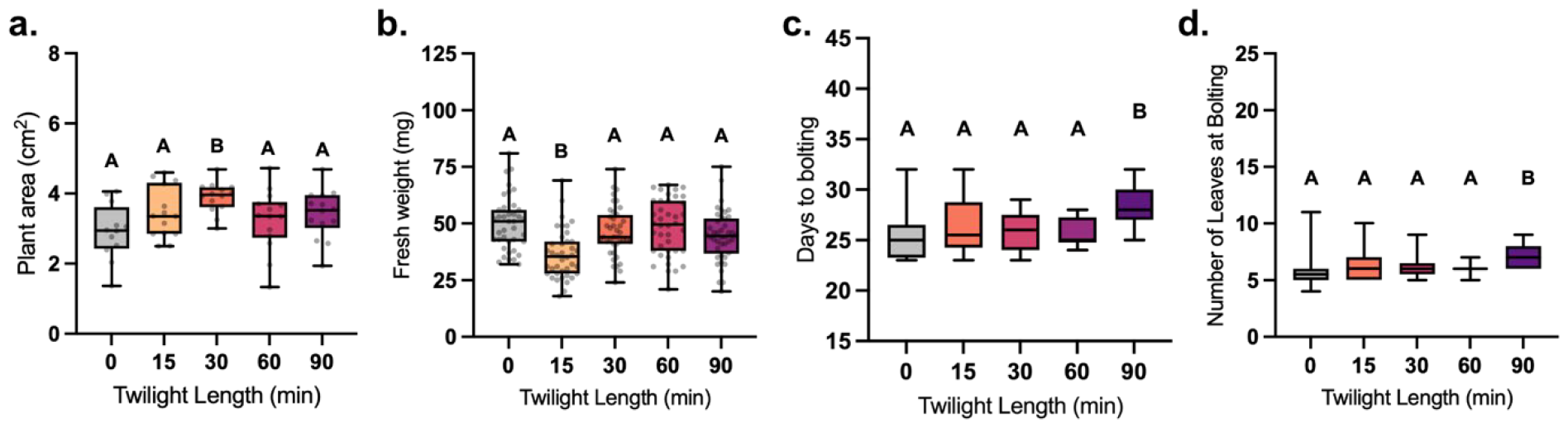
*lhy cca1* plants exhibit a muted phenotypic response to different twilight lengths. **(a**.) Leaf area of *lhy cca1* plants at 25 days post imbibition measured using PlantCV (n>12). **(b**.) Fresh weight measurements of *lhy cca1* plants at 25 days post imbibition. (n>39). **(c. & d**.) Flowering time measured in terms of **(c**.) number of leaves at bolting and **(d**.) days to bolting (n>22). Letters depict significantly different datapoints based on a one-way ANOVA and Tukey’s post-hoc test with adjusted p-value <0.05.

To identify the pathway from twilight length perception to the clock, we performed a screen of characterized homozygous photoreceptor knockout mutants under the 0 min, 30 min, and 90 min twilight length conditions. Here we found *phyd, phye* and *cry2* mutant lines most closely phenocopied the twilight-insensitive flowering response of *lhy cca1*, implicating these specific photoreceptors in twilight sensing (**Fig. S3**). Previous research has found connections between *PHYA, PHYB* and *CRY2* and photoperiodic flowering (*29–32*), as well as direct connections to the circadian clock through *LHY/CCA1* (*29, 33–37*).

Next, we sought to determine which of the flowering pathways was responsible for the twilight sensing phenotype. Hence, we examined flowering time mutants *gi-200, ft-10, soc1-2, flc-1* under different twilight conditions. *GI* is a key player in the photoperiodic flowering pathway (*11, 38, 39*), while *FLC* is a flowering regulator in autonomous flowering (*40*). *FT* encodes for the primary florigen signal to control flowering time (*41*), while *SOC1* is positively regulated by *FT* and negatively regulated by *FLC* to modulate flowering (*42*). Here, we found that *gi-200* and *ft-10* are unresponsive to twilight, phenocopying *lhy cca1*, while *soc1-2* and *flc-1* remain twilight responsive similar to wild-type plants (**Fig. S4**). Collectively, these data suggest that our twilight length dependent flowering phenotypes manifest via a *GI-FT* signaling mechanism, and not via a *FLC-SOC1* photoperiod-independent pathway.

To view the molecular impacts of increasing twilight duration in plants, we next performed a comparative quantitative proteomic and transcriptomic analysis of wild-type and *lhy cca1* plants. Plants grown under the same light conditions described above were sampled 25 days post imbibition, at ZT23, approaching the time of peak LHY/CCA1 expression. Proteomic analysis was carried out using our newly developed label-free BoxCarDIA quantitative proteomics methodology for more consistent protein quantification across the different twilight treatments (*43*). Transcriptomic analysis was performed using a modified Smart-seq2 protocol for cost-effective bulk RNA-sequencing (*44*). This allowed us to quantify a total of 5,748 protein groups and 21,126 transcripts, of which 4,746 protein groups and 11,728 transcripts were quantified across all the samples and replicates (**Supplementary Tables 7 & 8**). For both the protein and transcript datasets, we performed two differential analyses: (a) comparing WT and *lhy cca1* plants at each different twilight length (**Fig. 4 a & b; Supplementary Table 9**), and (b) comparing each twilight length to the no twilight treatment for each genotype (**Figure 4 c & d; Supplementary Table 10**). The former showed that the number of differentially abundant proteins in *lhy cca1* plants compared to wild-type increased under longer twilights. By contrast, the number of differentially expressed genes in *lhy cca1* plants peaked in the 30-min twilight treatment and was the lowest under 90 min of twilight. This pattern of transcriptional change appears to mirror phenotypic differences between *lhy cca1* and wild-type plants under different twilights, with both the transcriptome and phenotype of *lhy cca1* approaching that of wild-type plants under the extended twilight. We next focused on analysing proteins and transcripts changing across each twilight treatment compared to the no twilight condition for each genotype. Here, we identified much greater twilight-dependent protein-level regulation compared to transcriptome alteration in both genotypes. The number of differentially abundant proteins at each twilight treatment (compared to no twilight) increased linearly with twilight length for both genotypes (**Fig. 4c**) from 173 proteins in wild-type and 381 in *lhy cca1* under 15 min. of twilight to as many as 755 in wild-type and 722 in *lhy cca1* under the 90 min. twilight treatment. The number of differentially abundant proteins in common between the two genotypes also increased under longer twilights. In stark contrast, we found very few differentially expressed genes in both genotypes (but especially wild-type plants), with a maximum of 63 genes differentially expressed in wild-type plants grown under 15 min. of twilight (vs. no twilight) and 180 differentially expressed genes in *lhy cca1* plants grown under 60 min. of twilight (vs. no twilight) (**Fig. 4d**). Given that proteomics typically involves measurements of far fewer gene products than RNA-Seq analysis, it was surprising to find a greater number of differentially abundant proteins than differentially expressed genes. Hence, we believe the twilight response likely involves greater protein-level regulation (such as differential protein synthesis or degradation) than transcriptional regulation.

**Figure 4:**
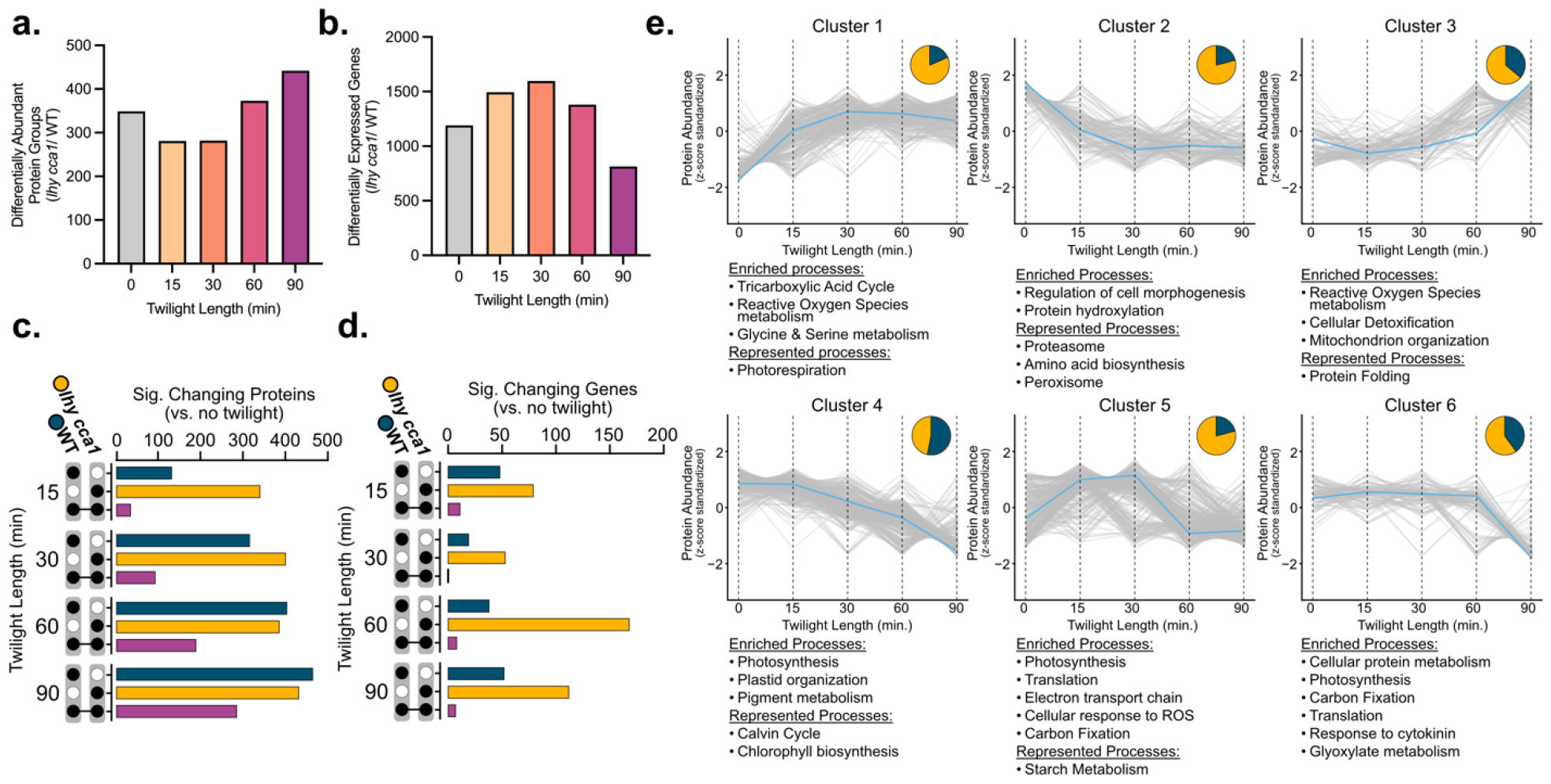
Proteome and transcriptome dynamics under different periods of twilight. **(a)** Number of proteins that change significantly in abundance at each of the different twilight durations in *lhy cca1* compared to wild-type plants. **(b)** Number of differentially expressed genes between *lhy cca1* and wild-type plants at each twilight length. **(c)** Upset plot showing the overlaps in proteins significantly changing in abundance and **(d)** differentially expressed genes under different twilight durations (compared to no twilight) in both *lhy cca1* and wild-type plants. **(e)** Shape-based clustering of protein abundance (z-score standardized) with clusters annotated with enriched GO-terms and selected member proteins and pathways. Blue lines show cluster centroids. Pie charts show proportion of proteins in each cluster found in *lhy cca1* (yellow) and wild-type (blue) samples.

To identify biological processes potentially subject to such protein-level regulation, we next clustered proteins quantified across all replicates and treatments in our analysis by the shape of their z-score standardized abundance over different twilight lengths. Clustering analysis was performed on the subset of proteins that changed significantly in their abundance dependent on twilight length (ANOVA p-value < 0.05) in each genotype. We partitioned the data into 6 clusters depending on the shape of their abundance trajectories, thereby clustering proteins whose abundance changed similarly over different twilight lengths (**Fig. 4e**). We then performed gene ontology enrichment analysis for each protein cluster as well as StringDB analysis (*45*) to identify both enriched biological processes as well as other pathways represented by the constituent proteins in each cluster. Here, we found that clusters 1, 2, and 5 were dominated by proteins changing in abundance in *lhy cca1* plants, while clusters 3, 4, and 6 contained a higher number of proteins changing in abundance in wild-type plants. Overall, proteins in clusters 1, 3, and 5 followed trajectories that either paralleled or anti-paralleled the growth phenotypes observed, either peaking or reaching their lowest abundance under 30-60 min of twilight. Interestingly, the two clusters that contained proteins increasing in abundance over longer twilights (1 and 3) were enriched in proteins relating to reactive oxygen species metabolism, glycine & serine metabolism, and cellular detoxification. Cluster 1 also contained proteins involved in photorespiration (which intersects with glycine and serine metabolism). This suggests the hypothesis that optimal lengths of twilight might allow plants to more appropriately manage photooxidative stress. We also found proteins involved in proteasome and amino acid biosynthesis to be enriched in cluster 2 and proteins involved in cellular protein metabolism enriched in cluster 6, perhaps explaining the difference between the number of differentially abundant proteins and transcripts observed in **Fig. 4 (c & d)**. Finally, we also found proteins involved in photosynthesis, both in light harvesting and carbon fixation, to be enriched in clusters 4, 5, and 6. Overall our -omics results highlight wide-ranging proteome-level changes associated with longer twilight lengths, specifically implicating management of oxidative stress and photosynthesis in conjunction with proteasome and translational regulation in the observed twilight-responsive phenotypic changes in wild-type and *lhy cca1* plants.

## Conclusions

A central question in circadian biology is: “How do plants measure the length of day and night across changing seasons?”. The use of rectangular LD cycles to perform photoperiod experiments in the past has implied a simplified go/no-go process whereby plants measure daylength simply by timing the number of hours between the instantaneous onset of light from one day to the next. This view would suggest that plants perceive day and night as a digital “on-off” signal rather than the gradual transition it is in real life. By comparing rectangular LD photoperiods and a series of more natural twilight ramps, we show that the plant clock (via its LHY/CCA1 module) affects development not simply by measuring the total number of hours or minutes of light, but via more subtle tuning by the *amount of light incident over a time interval* at the beginning and end of the day (i.e., the slope of light intensity over time).

Another alternative hypothesis for daylength perception remains that plants might perceive gradual changes in light intensity in the mornings and evenings but nevertheless convert this analog signal to a binary output when triggering photoperiodic flowering, by imposing a high irradiance threshold, similar to that recently discovered for morning *FT* induction via *PHYA* (*17*). Our findings do not entirely contradict this idea, but the results depicted in Fig. 2 impose a threshold of at least 40 μmol m^−2^ s^−1^ for such a high-irradiance threshold model. Further, our mutant screen does not implicate *PhyA* in twilight length sensing. Our follow-up genetic analyses (along with the published literature) allow us to postulate a linear genetic pathway from *PHYD, PHYE*, to *LHY/CCA1* to *GI* and *FT* as being responsible for the twilight length-dependent flowering phenotype we observed (**Figure 5**).

**Figure 5:**
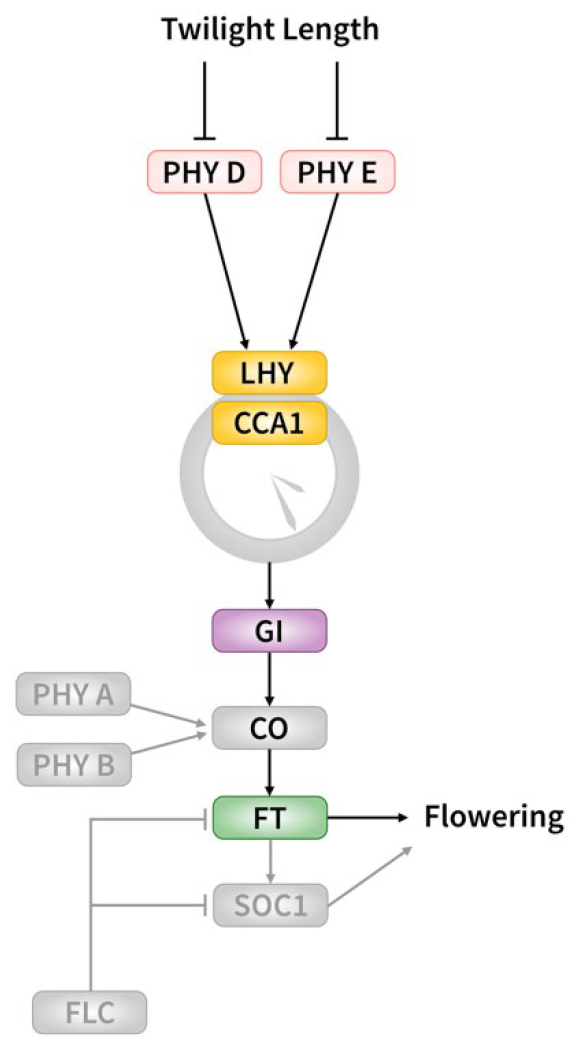
Proposed genetic pathway connecting twilight length to flowering time. Genes depicted color are genetically linked to twilight length sensing according to this study while greyed out genes are not.

Future work will aim to study this proposed pathway in greater depth and delineate the roles of these genes in twilight vs. laboratory-based rectangular photoperiod sensing. Further, our combined transcriptomic and proteomic analysis leads us to hypothesize that the growth and developmental phenotypes impacted by changing twilight lengths are the result of systemic proteome-level regulation (via a regulatory proteasomal-translational control axis) of reactive oxygen species metabolism, photosynthesis, and carbon metabolism. Future targeted studies may resolve exactly how clock control of twilight perception alters these processes to optimize growth during natural photoperiod transitions.

Collectively, we believe our results offer a more refined understanding of how the circadian clock interprets daylength changes in natural environments. It should be noted that our experimental design focused exclusively on studying the impact of twilight length by controlling the daily light integral (DLI) between treatments. Natural twilight length variation is of course not independent of changes in DLI and includes changes in spectral composition over the course of a day. Future work in this domain could iterate our experimental design to factor in more realistic twilight and spectral changes, including at different latitudes, different altitudes, and the impact of changes in cloud cover or pollution among other geographical influences on plant growth and development. Furthermore, given the molecular complexity and agronomic value of the traits measured in this study (i.e., growth and flowering time), we expect our systems-level molecular results to inform future targeted work focused on identifying genes and pathways influencing photoperiodic flowering and diel plant growth modulation over natural light transitions, leading to new breeding and engineering targets. In conclusion, our results add to a growing focus on understanding plant chronobiology from a biogeographical point of view (*46*), and demonstrate that simulating more natural environmental changes can lead to a more nuanced understanding of well-studied clock controlled processes, such as photoperiodism.

## Methods

### Growth cabinet design and construction

Growth cabinets with ventilation holes were constructed using black acrylic and housed in Conviron walk-in growth chambers. The inner walls of the cabinets were lined with reflective Mylar material to ensure uniform lighting. Light bars were equipped with 6 different LEDs (G2V Optics Inc. Alberta, Canada; Wavelength details provided in **Supplementary Table 1**) programmed to approximate the AM1.5 reference spectrum.

### Plant growth

*Arabidopsis thaliana* (Col-0), *lhy-20 cca1-1* (*lhy cca1*; (*47*)), *rve 4-1 6-1 8-1* (*rve 4 6 8*; (*48*), *elf4-101* (*49, 50*) and *toc1-2* (*51*), along with *gi-200* (26), *soc1-2*(*52*), *ft-10*, and *flc-1* (Arabidopsis Biological Resource Center) seeds were sown on 0.5x MS agar and stratified for 3 days and then germinated in separate growth cabinets, each with a different light regime. After 4 days, seedlings were transplanted into pots filled with soil. Homozygous photoreceptor mutants *phyA, phyB, phyC, phyD, phyE, cry1, cry2* and *cry1 cry2* plants (*53*) were stratified, germinated and grown as described above. For imaging and flowering time experiments, one seedling was grown per pot (in 2.5-inch pots). For proteomics and RNA-seq, six seedlings were grown in larger 4-inch pots.

### Imaging and computer vision analysis

Plants were imaged using Arducam 5MP OV5647 cameras (https://www.arducam.com/product/arducam-m12-night-vision-ir-cut-raspberry-pi-camera/), with images taken every 5 min between ZT0 and ZT12 for multiple days. Images were then processed using a customized version of PlantCV to identify individual plants from a whole tray image, measuring their total leaf area and perimeter. Briefly, one or more representative images from each growth chamber configure our PlantCV pipeline and output a configuration file that is used to bulk analyze images taken by the same camera for each experiment. These images are first undistorted using the camera intrinsic matrix and distortion coefficients obtained by calibrating cameras with the OpenCV fisheye camera model. The white balance of the images is then adjusted using a reference white spot fixed on our plant growth trays and the built-in PlantCV ‘white_balance’ function. Regions of interest (ROIs) around each individual plant are then manually drawn using a custom graphical user interface (GUI) ROI Drawing Tool that was based on a tool created by Github user BoKuan Liu (https://github.com/DennisLiu1993/Zoom-In-Out-with-OpenCV). Next, images are segmented into ‘plant’ and ‘background’ with a colour threshold using a custom GUI based HSV Thresholding Tool. After successful thresholding, optimized parameters are exported into a JSON configuration file. Subsequently, all images obtained from each camera for an experiment are analysed using the configuration file produced for that camera. The image processing pipeline accepts the configuration file as input and produces two types of outputs. The first output is a comma-separated values (CSV) file for each camera, containing extracted plant traits along with metadata for each sample. The second output is the set of annotated images produced by PlantCV, which can be used to visually inspect the quality of the image analysis. Final plant area analysis across twilight lengths was performed by averaging the area of each plant measured during a 1-hour period (ZT 5.5-6.5) at 25 days post imbibition to avoid interference from leaf movement effects. Next, the averaged area at 25 days post imbibition per plant was plotted and a one-way ANOVA and Tukey’s post-hoc test with an adjusted p-value threshold of 0.05 was applied to detect significant changes between twilight treatments. Final plant area measurements are provided in Supplementary Table 2. Raw and processed images can be downloaded from www.doi.org/10.5281/zenodo.7977125.

### RNA-Seq and transcriptome data analysis

Total RNA was extracted from rosette tissue harvested at ZT 23 on day 25 post imbibition (n = 4 biological replicates) using the Direct-zol-96 MagBead RNA kit (Zymo Research). RNA was checked for degradation using a Tapestation 4150 (Agilent) following the manufacturer’s protocol. Cost-effective RNA-Seq was performed using a miniaturized library preparation protocol by applying the Smart-seq2 single-cell method on bulk RNA. For this, cDNA was generated following the protocol as described in Picelli et al. (*44*) with the following adjustment: 1 μl of lysate was transferred from the lysis plate into a new plate containing the RT mix. Preamplification of cDNA used 23 PCR cycles and was purified using Agencourt Ampure XP beads (Beckman Coulter) with a modified bead: DNA ratio of 0.8x. The quality of cDNA was checked using a NGS Fragment High Sensitivity Analysis Kit (Advanced Analytical) and a Fragment Analyzer (Advanced Analytical). The cDNA concentration was measured using a qubit High sensitivity dsDNA Kit. Libraries were prepared using a Nextera XT DNA Library Preparation Kit (Illumina), using a standard protocol but with all reaction volumes reduced by 1/10 to accommodate the automation of the prep on the Echo LabCyte liquid handler (Beckman). Libraries were purified using Agencourt Ampure XP beads (Beckman Coulter). Size distribution of library pools was checked using a Fragment Analyzer and a NGS Fragment High Sensitivity Analysis Kit. Samples were pooled equimolar, and the final pool quantified with the Kapa library quantification kit (Roche). The final pool was sequenced on a NovaSeq 6000 to produce paired-end 150 bp reads, with an average read depth of 10M reads per sample.

Protocol Quality control of raw reads was performed with FastQC ver. 0.11.7 (*54*). Adapters were filtered with Trimmomatic v0.39 (*55*) . Splice-aware alignment was performed with Star (*56*) against the Arabidopsis Araport 11 reference genome using the default parameters. Reads mapping to multiple loci in the reference genome were discarded. Quantification of reads per gene was performed with FeatureCounts from Subread package (*57*). Count-based differential expression analysis was done with the R-based Bioconductor package DESeq2 (*58*). Reported p-values were adjusted for multiple testing by controlling the false discovery rate using the Benjamini-Hochberg procedure; Log2 Fold-Change > 0.58.

### Proteomics sample preparation, mass-spectrometry, and data analysis

#### Tissue Extraction and Sample Processing

Frozen rosette tissue (n=4 biological replicates) was ground using Geno/Grinder (SPEX SamplePrep) for 30s at 1200 rpm and aliquoted into 100 mg fractions under liquid N2 prior to extraction. Ground tissue was then dissolved in protein extraction buffer (50 mM HEPES-KOH pH 8.0, 100 mM NaCl, and 4% (w/v) SDS at a 1:3 (w/v) ratio and extracted by shaking 1000 RPM 95 °C for using a tabletop shaker (Eppendorf ThermoMixer F2.0). Samples were then centrifuged at 20,000 xg for 10 min at room temperature and the supernatant retained in new tubes. Protein extracts were then reduced with 10 mM dithiothreitol (D9779, Sigma) for 5 min at 95 °C, followed by alkylation for 30 min with 55 mM Iodoacetamide (I1149, Sigma) at room temperature. Total proteome peptide fractions were then generated using a Kingfisher Apex (ThermoFisher Scientific) automated sample processing system as outlined by Leutert et al. (2019) without deviation (*59*). Peptides were digested using sequencing grade trypsin (V5113; Promega) diluted in 50 mM Triethylammonium bicarbonate buffer pH 8.5 (T7408; Sigma). Following digestion, samples were acidified with trifluoroacetic acid (A117, Fisher) to a final concentration of 0.5% (v/v). Peptides were desalted as previously described (*60*) using an OT-2 liquid handling robot (Opentrons Labworks Inc.) mounted with Omix C18 pipette tips (A5700310K; Agilent). Desalted peptides were dried and stored at - 80°C prior to re-suspension in 3.0% (v/v) ACN / 0.1% (v/v) FA prior to MS injection.

#### LC-MS/MS Analysis

Peptides were analyzed on a Fusion Lumos Orbitrap mass spectrometer (ThermoFisher Scientific) in data independent acquisition (DIA) mode. A total of 1 μg of re-suspended peptide was injected per replicate using and Easy-nLC 1200 system (LC140; Thermo Scientific) mounted with an Acclaim PepMap 100 C18 trap column (Cat# 164750; ThermoFisher Scientific) and a 50 cm Easy-Spray PepMap C18 analytical column (ES903; ThermoFisher Scientific) warmed to 50 °C. Peptides were eluted at 300 nL/min using a segmented solvent B gradient of 0.1% (v/v) FA in 80% (v/v) ACN (A998, Fisher) from 4 to 41% solvent B (0 – 120 min). A positive ion spray voltage of 2.3 kV was used with an ion transfer tube temperature of 300 °C and an RF lens setting of 40%. BoxCar DIA acquisition was also performed as described (*43*) without any deviation. Briefly, MS1 spectra were acquired using two multiplexed targeted SIM scans of 10 BoxCar windows each. Full scan MS1 spectra (350–1400 m/z) were acquired with a resolution of 120 000 at 200 m/z and normalized AGC targets of 100% per BoxCar isolation window. Fragment spectra were acquired a resolution of 30 000 across twenty-eight 38.5 m/z windows overlapping of 1 m/z using a dynamic maximum injection time and an AGC target value of 2000%, with a minimum number of desired points across each peak set to 6. HCD fragmentation was performed using a fixed 27% fragmentation energy.

#### Data analysis

Data was analysed using Spectronaut ver. 17 (Biognosys AG), with all analysis details found in the in the Spectronaut result summary (Supplementary Table 2). Significantly changing differentially abundant proteins were determined and corrected for multiple comparisons (Bonferroni-corrected p-value <0.05; q-value; Log2 Fold-Change; Log2FC >0.58) (**Fig. 4 a-d**), while significantly changing proteins plotted across each time-point (**Fig. 4e**) were deduced using Perseus ver. 1.6.15.0 (*61*), with row valid values set to 100% and a Benjamini Hochberg-corrected ANOVA p-value <0.05.

## Acknowledgements

The authors would like to thank Prof. Dr. Jose Pruneda-Paz (University of California - San Diego) for providing *lhy-20 cca1-1* seeds and Prof. Dr. Stacey Harmer (University of California – Davis) for providing *rve 4-1 6-1 8-1* and *toc1-2* seeds. The authors acknowledge G2V Optics Inc. for lighting equipment and support, Dr. Jack Moore from the Alberta Proteomics and Mass-Spectrometry Facility, and Annelien Verfaillie, Khaled Mirzai, and Alvaro Cortes Calabuig from the GenomicsCore Leuven for technical support.

## Author Contributions

DM: Conceptualization; Investigation; Formal Analysis; Methodology; Data curation; Writing original draft; Visualisation.

SS: Investigation; Methodology; Data curation; Review & Editing MT: Investigation; Formal Analysis; Review & Editing

CK: Software; Data curation; Review & Editing MR, LG: Investigation

ES: Formal Analysis; Review & Editing

RGU: Conceptualization; Methodology; Supervision; Project administration; Data curation; Writing original draft; Review & Editing; Funding acquisition.

## Data availability

All raw proteomics data can be found through the PRoteomics IDEntifications Database (PRIDE) using the dataset identifier PXD039428. All raw image data can be found on Zenodo (www.doi.org/10.5281/zenodo.7977125). RNA sequencing data has been deposited in the European Nucleotide Archive (ENA) at EMBL-EBI under accession number PRJEB62130. Code used to analyze image data, perform k-means clustering, and plot graphs is available on Github at: https://github.com/UhrigLab/twilight_length

## Supplementary Materials

Figure S1-S4 Tables S1 to S9

1. Wavelength range of LEDs and ramping program
2. Leaf area measurements and statistical analysis
3. Fresh weight measurements and statistical analysis
4. Flowering time measurements and statistical analysis
5. Flowering time measurement under rectangular photoperiods and statistical analysis
6. Proteome quantification data
7. Normalised counts from RNA-Seq analysis
8. Candidate genes and proteins in WT plants compared to *lhy cca1*
9. Candidate genes and proteins per twilight length in each genotype compared to no twilight treatment

## Supplementary Figures

**Figure S1:**
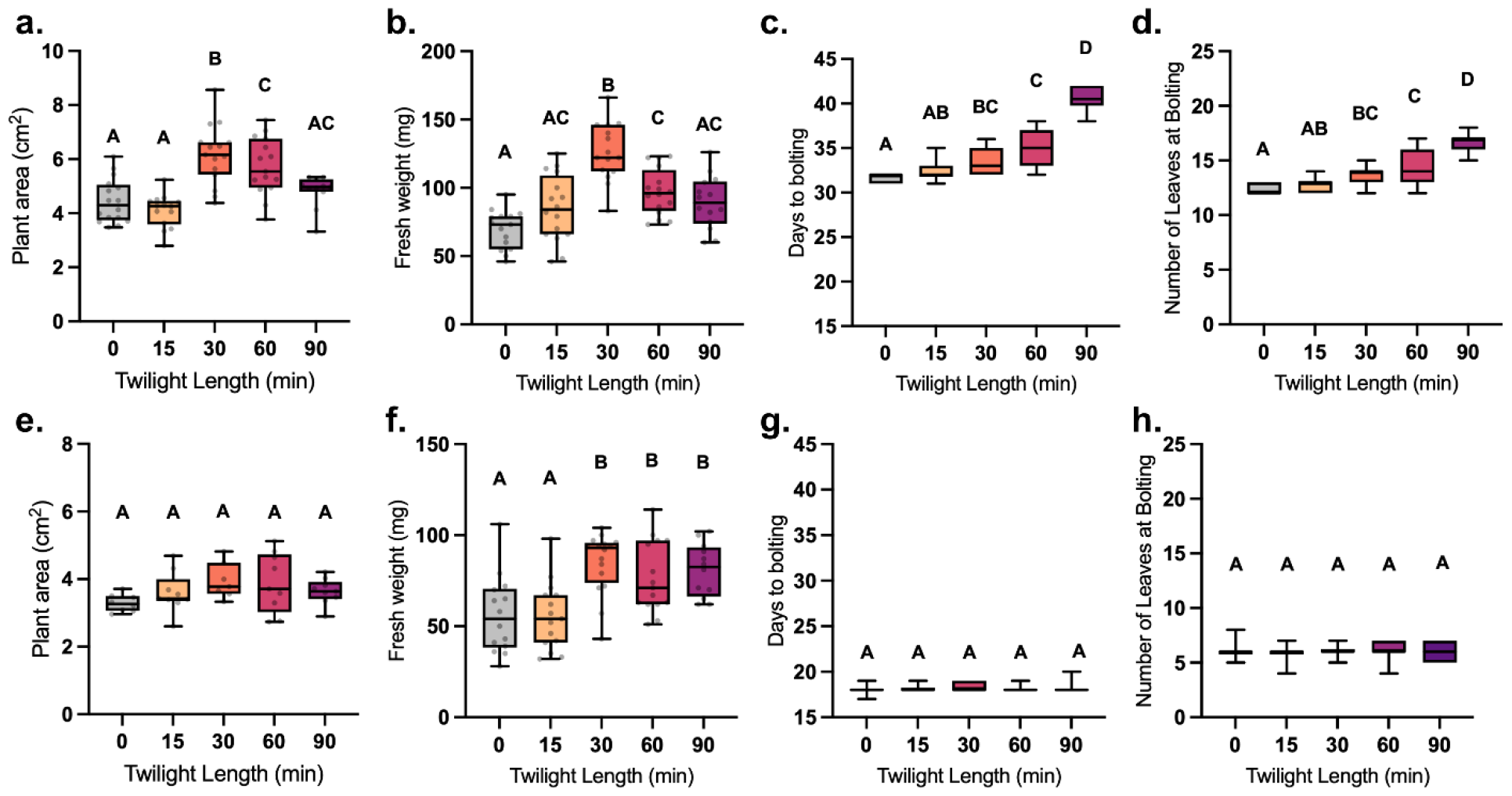
Repetition of Col-0 (a-d) and *lhy cca1* (e-h) twilight phenotypic data (n>8). Letters depict significantly different data points based on a one-way ANOVA and Tukey’s post-hoc test with adjusted p-value <0.05.

**Figure S2:**
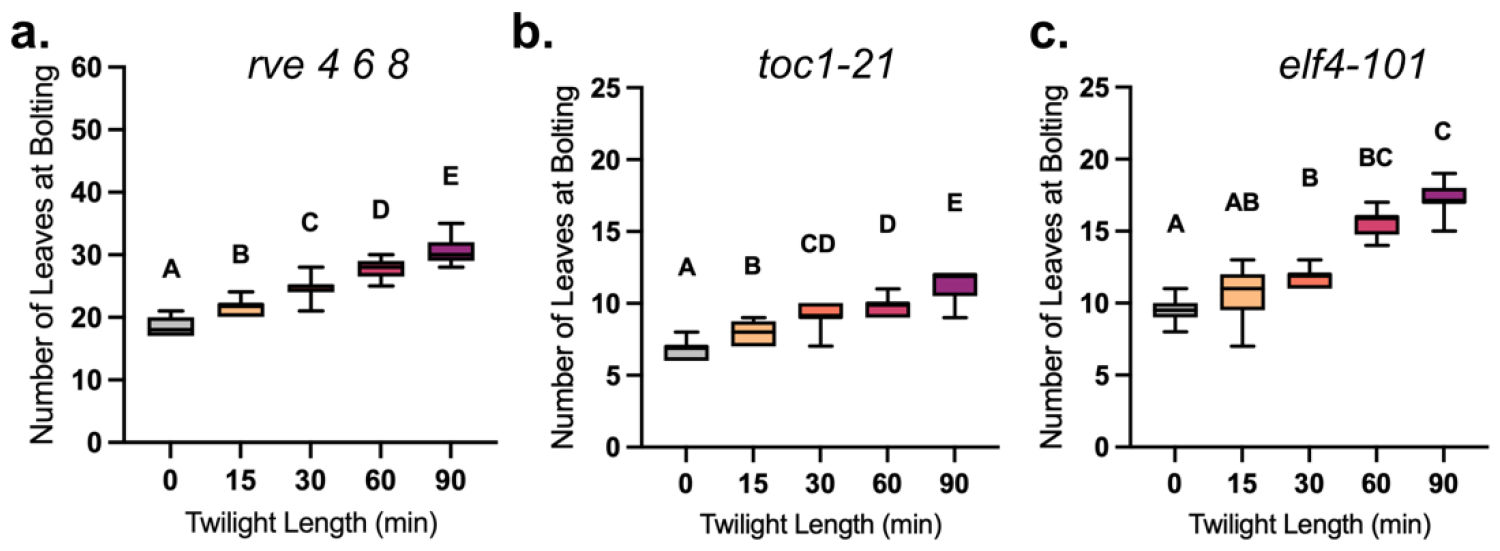
Flowering time measured in terms of number of leaves at bolting for clock mutants *rve 4 6 8, toc1-21* and *elf4-101* (n>20). Letters depict significantly different data points based on a one-way ANOVA and Tukey’s post-hoc test with adjusted p-value <0.05.

**Figure S3:**
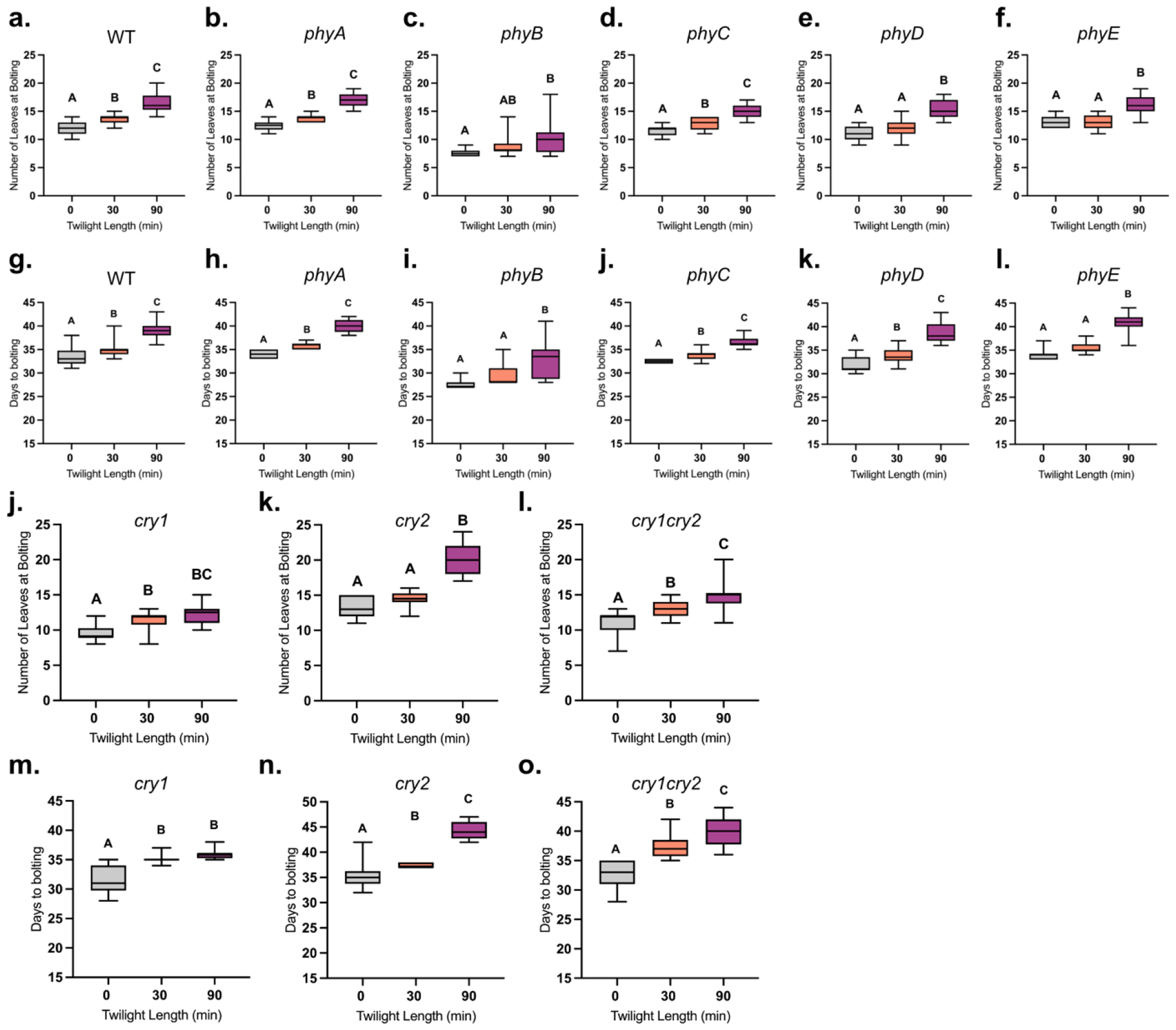
Flowering time measured in terms of number of leaves at bolting (a-f, j-k) and days to bolting (g-l; m-o) for single phytochrome (*phya – e*) and cryptochrome (*cry1, cry2* and *cry1 cry2)* mutant plants (n>14). Letters depict significantly different data points based on a one-way ANOVA and Tukey’s post-hoc test with adjusted p-value <0.05.

**Figure S4:**
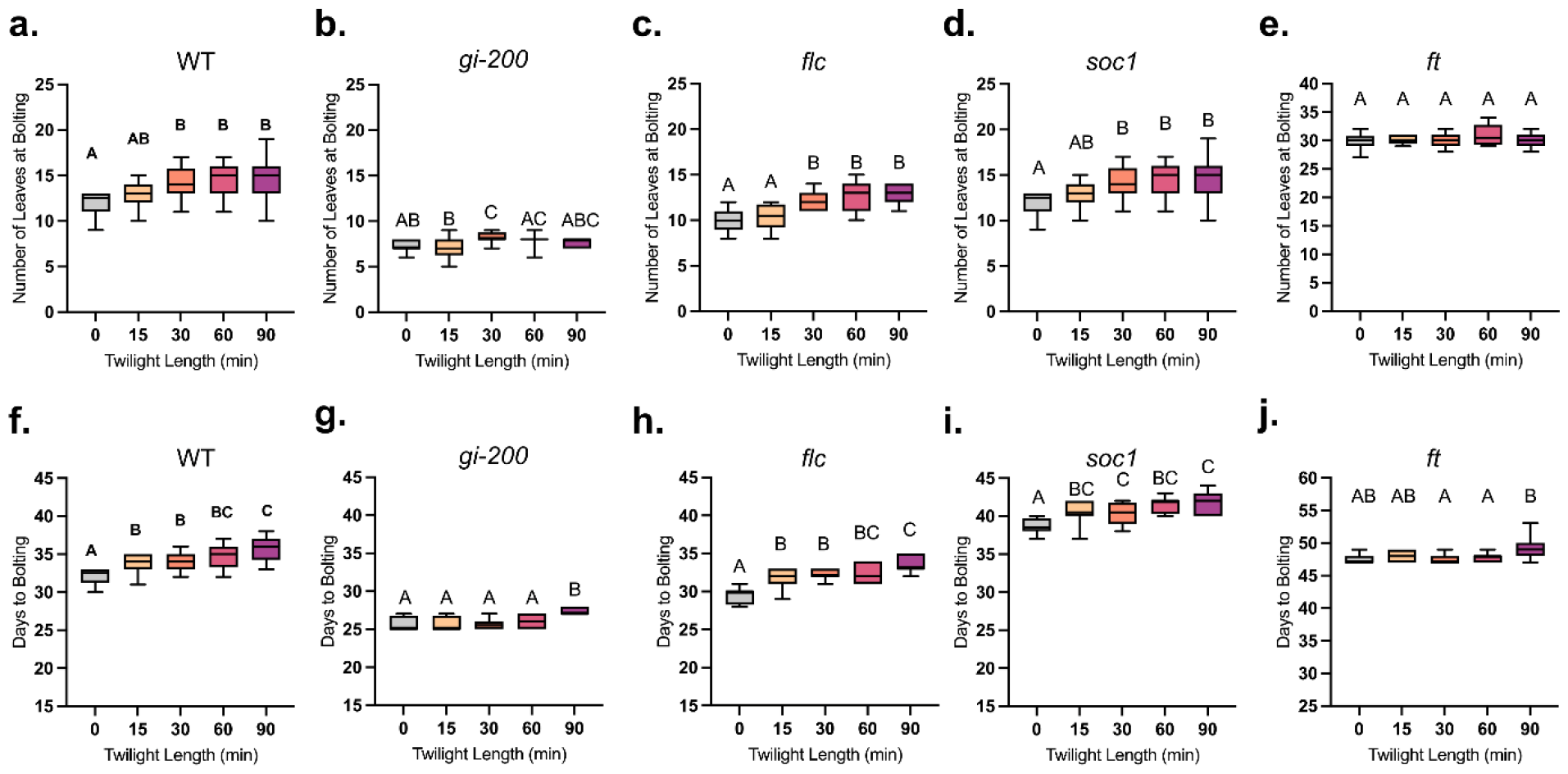
Flowering time measured in terms of number of leaves at bolting (a-e) and days to bolting (f-j) for flowering related mutants (n>8). Letters depict significantly different data points based on a one-way ANOVA and Tukey’s post-hoc test with adjusted p-value <0.05.

